# Characteristics of the rumen virome in Japanese cattle

**DOI:** 10.1101/2023.03.20.532305

**Authors:** Yoshiaki Sato, Hiroaki Takebe, Kento Tominaga, Jumpei Yasuda, Hajime Kumagai, Hiroyuki Hirooka, Takashi Yoshida

## Abstract

The rumen microbiome is a highly complex ecosystem that includes bacteria, archaea, protozoa, fungi, and viruses. Viruses have a high potential to modify the rumen digestion of feeds via infection and cell lysis of prokaryotes in the rumen; however, understanding of the rumen virome is substantially less advanced due to limitations of the reference genome database. In this study, we conducted metagenomic sequencing of virus-like particles (VLPs) in the rumens of 22 Japanese cattle to construct a reference viral genome catalog of the rumen and uncover the rumen virome characteristics. We succeeded in construction of 8 232 nonredundant viral genomes (≥5 kb length and ≥50% completeness). Among them, putative hosts of 1 223 virus genomes were predicted, and 1 053 virus genomes were taxonomically classified, mainly Siphoviridae, Myoviridae, and Podoviridae. Additionally, 2 764 putative auxiliary metabolic genes (AMGs) were identified in the viral genomes. Importantly, 22 viral genomes associated with archaea in the rumen were identified. Some archaeal viruses have AMGs related to DNA synthesis, suggesting that archaeal viruses control archaeal populations in the rumen and affect methane production from the rumen. Furthermore, we revealed that most rumen viruses were highly rumen-and individual-specific and related to rumen-specific prokaryotes. Overall, the rumen viral catalog and findings of this study will help future analyses to uncover the roles of rumen viruses in feed digestion, productivity, and methane production.

## Introduction

Ruminants have evolved into four chambers: the rumen, reticulum, omasum, and abomasum. The rumen, the first chamber of ruminants, is a large fermentation chamber for feed ingested by the rumen microbiome. The rumen microbiome is a complex ecosystem comprised of bacteria, protozoa, fungi, archaea, and viruses [1, 2]. The microbiome in the rumen, mainly bacteria, breakdowns, and ferments plant materials, such as cellulose and hemicellulose, producing volatile fatty acids as the primary energy source for host ruminants. Thus, ruminants can convert human-indigestible feed to human-digestible food, such as milk and meat, owing to the rumen microbiome.

Viruses are estimated to be abundant in the rumen, ranging from 5 × 10^7^ to 1.6 × 10^10^ particles/mL of ruminal fluid [3, 4]. Among them, bacteriophage (hereafter “phage”), which infects bacteria, has been the most well-characterized in the rumen, although there are also viruses that can infect archaea, fungi, and protozoa [5]. Viruses may play crucial roles in the rumen ecosystem via infection and cell lysis of bacteria, reproduction, and reprogramming of microbial metabolism [6, 7, 8, 9]. Thus, it is expected that viruses in the rumen have a high potential of indirectly modifying the rumen digestion and fermentation of feeds, hence animal performance and methane production, by changing the rumen microbial ecology.

With the development of high-throughput sequencing technology, culture-independent metagenomics has better revealed the viral community (virome), which enables the identification of uncultured DNA viruses. Owing to the lack of a universal marker gene across viral genomes, shotgun metagenomic sequencing is needed to fully address viral communities. Despite the limitations of the methodology, a few studies have been performed using metagenomic technology for rumen samples, and they provide fundamental knowledge of the rumen virome [6, 7, 8, 10]. For example, Berg Miller *et al.* [6] demonstrated that abundant viral families in the rumen were Myoviridae, Siphoviridae, Mimiviridae, and Podoviridae, and most viruses were related to the dominant bacteria in the rumen, such as Firmicutes, Proteobacteria, and Bacteroidetes. A recent study reported that virus-encoded auxiliary metabolic genes (AMGs), which reprogram the microbial host metabolism, may facilitate the breakdown of complex carbohydrates and augment energy production in the rumen [7]. However, understanding of the rumen virome is substantially less advanced than that of the community of prokaryotes, and the impact of viruses has been overlooked in most studies on the rumen microbiome.

For metagenomic analysis of virome, reference viral genomes are important because they are used for taxonomic classification, estimation of viral taxa distribution, and identification of host-virus interactions and functional potential [11]. To date, many metagenomic sequencing studies have been conducted to construct viral genomes in other environments. For example, more than 33 200 [12], 142 000 [13], and 189 600 viral genomes [14] have been identified in the human gut. Additionally, the IMG/VR database, the largest collection of publicly available viral genomes, primarily comprises genomes derived from the marine, freshwater, and human gut [11]. However, a reference viral genome database for the rumen has not yet been constructed. Considering that the majority of viruses have habitat-type specificity [15], a rumen viral genome catalog is required to enable metagenomic analysis of the rumen virome. Additionally, to understand how the rumen virus influences the rumen microbiome, it is important to identify the host-phage interactions and AMGs harbored within the viral genomes.

In the present study, to generate and characterize the rumen virus genome catalog, we conducted metagenomic sequencing of virus-like particles (VLPs) from the rumen of 22 Japanese cattle. As a result, 8 232 nonredundant viral genomes with ≥5 kb length and ≥50% completeness were described, vastly expanding the diversity of rumen viruses. Notably, 22 viral genomes related to archaea, which may be associated with methane production in the rumen, were identified. Additionally, we revealed that most rumen viruses were highly rumen-and individual-specific and were related to rumen-specific prokaryotes.

## Materials and methods

The experimental design and protocol were approved by the Kyoto University Animal Ethics Committee (permit no. R2-119).

### Animals and rumen sampling

The experimental animals and designs have been described previously [16, 17]. Briefly, rumen samples were collected from Japanese Black (JB; n = 8), Japanese Shorthorn (JS; n = 2), and Japanese Black sires × Holstein dams crossbred (F1; n = 12) steers using stomach tubing. Among them, six JB and six F1 steers were fed an identical diet and kept in a single cattle barn [16]. The rumen samples were filtered through four layers of cheesecloth, transported on dry ice to our laboratory, and preserved at −80 °C until use.

### Virus-like particles enrichment, DNA extraction, and metagenomic sequencing

For VLP isolation, rumen samples were thawed and centrifuged at 6,000 × *g* for 30 min to remove large debris. The supernatants were passed through a 0.45 μm followed by 0.22 μm PVDF filter (Millipore, Burlington, MA, USA) to remove residual host and bacterial cells. The number of VLPs were counted by diluting the filtered samples to × 10^4^ or × 10^5^ in virus-free water, which was 0.02 μm-filtered MilliQ water. The VLPs in 1 mL of the diluted samples were strained using SYBR-gold for 20 min and captured on a 0.02 μm Anodisc filter (Whatman, Buckinghamshire, UK). After washing twice with 1 mL of virus-free water, VLPs on the filter were visualized using an epifluorescent microscope. For VLP purification, CsCl density-gradient centrifugation was performed as described by Hurwitz *et al*. [18], with minor modifications. Briefly, filtered samples were deposited onto CsCl density gradients composed of 1.7, 1.5, and 1.45 g/mL layers and centrifuged for 1 h at 28,500 rpm at 15 °C using an Optima XE-90 Ultracentrifuge and SW 41 Ti Swinging-Bucket Rotor (Beckman Coulter, Inc., Pasadena, CA, USA). After centrifugation, we retrieved the 1.45–1.5 g/mL layer using a sterile syringe and needle. The purified VLPs were dialyzed in SM buffer (100 mM NaCl, 50 mM Tris-HCl, pH 7.5, 8 mM MgSO_4_) using 30 kDa super-filters. The purified VLP fraction was treated with DNase I (Ambion) at 37 °C for 1 h. DNA extraction was performed using the previously described xanthogenate method [19]. The extracted DNA was used for library preparation using the Nextera XT DNA library preparation kit (Illumina, San Diego, CA, USA). Libraries were sequenced using Illumina HiSeq X Ten (2 × 150-bp).

### Bioinformatic analysis

#### Assembly and viral genome identification

Low-quality reads and Illumina adapters were trimmed using Trimmomatic v0.39 [20]. After filtering, 1.05 G reads were retained, with 47.94 ± 11.1 M reads per sample. The filtered reads of each sample were assembled with SPAdes v3.15.3 using the “--meta” option [21, 22]. The assembled contigs longer than 5 kb were retained. All retained contigs were clustered at a 95% identity and 85% coverage using CD-HIT v4.8.1 [23]. Nonredundant contigs were screened using VirSorter2 v2.2.2 [24]. We defined nonredundant viral contigs, classified as viral candidates by Virsortor2 [24], as viral operational taxonomic units (vOTUs). Subsequently, the completeness of vOTUs was assessed using CheckV v0.8.1 [25]. Additionally, CheckV [25] was used to assess host-virus boundaries and to remove the host fraction from vOTUs. The vOTUs that were assigned to complete high-quality (≥90% completeness) or medium-quality (50–90% completeness) were collected (hereafter defined as RVG1) and used for further analysis.

#### Taxonomic classification and viral protein cluster determination

We constructed a dataset of rumen viral genomes to compare with other public ruminal viral contigs. First, the raw sequencing data of the deep rumen viral metagenome [7] were downloaded from the Sequence Read Archive under accession numbers SRR3621238, SRR3621239, SRR3621240, SRR3621241, SRR3621242, and SRR3621244. The data were trimmed and assembled in the same manner as described above. The assembled contigs were combined with moose viral contigs [8] downloaded from the JGI GOLD database (https://gold.jgi.doe.gov/) corresponding to Ga0207349, Ga0207350, Ga0207351, and Ga0207352. We also performed CD-HIT [23], VirSorter2 [24], and CheckV [25] on contigs longer than 5 kb, as mentioned above. The rumen viral contigs with ≥50% completeness (n = 979) were retained (hereafter defined as RVG2) and combined with RVG1 for further analysis using vConTACT2 [26].

Proteins were predicted with Prodigal v.2.6.3 [27] with “-p meta,” and the predicted protein sequences were treated as the input for vConTACT2. vConTACT2 v.0.9.19 [26] was performed with its ProkaryoticViralRefSeq201-Merged database using all-vs-all DIAMOND protein comparison, MCL for protein clustering, and ClusterONE for genome clustering. The resulting gene-sharing network was visualized using Cytoscape v.3.9.1 [28]. Taxonomy at the family level was assigned to RVG1 vOTUs that shared a viral cluster (VC) with one or more RefSeq genomes. RVG1 vOTUs that were grouped into ambiguous clusters (i.e., outliers and overlap) were identified as singletons. For the RVG1 vOTUs that were taxonomically unassigned by vConTACT2, the genome similarity score (*S_G_* values) derived from the all-against-all tBLASTx computation was calculated using ViPTree with the default prokaryotic viral genomes [29]. RVG1 vOTUs with *S_G_* > 0.15 against complete viral genomes in the database were taxonomically assigned at the family level. Next, we manually checked the results of vConTACT2 and ViPTree. Taxonomy was annotated to RVG1 vOTUs that shared a VC with one or more RVG1 vOTUs taxonomically assigned by ViPTree. RVG1 vOTUs in a VC assigned to more than two families were defined as an “unclassified family.” ViPTree [29] was also used to construct a proteomic tree for archaeal viruses reported in this study and the Virus-Host DB [30].

The predicted protein sequences were mapped to the Pfam database [31] using HMMER v.3.3 [32], with an E value < 10^−5^. RVG1 vOTUs with genes encoding integrase or recombinase (PF00239, PF00589, PF02899, PF07508, PF09003, PF09299, PF10136, PF13356, PF14659, PF16795, and PF18644) were classified as lysogenic viruses.

#### Auxiliary metabolic genes identification

To be conservative, RVG1 vOTUs classified as proviruses via CheckV [25] were excluded for identification of *bona fide* viral AMGs. Before the analysis, AMGs and VirSorter2 [24] were performed again with the “--prep-for-dramv” option against the filtered RVG1 vOTUs (n = 8 181) to generate “affi-contigs. tab” files. Putative AMGs were annotated using the DRAM-v [32]. For the DRAM-v output, genes with an auxiliary score < 4,-M, and-F flags were regarded as putative AMGs. Genome linear maps of archaeal RVG1 vOTUs with AMGs were visualized using Easyfig 2.2.2 [34].

#### Virus-host predictions

Microbial hosts for RVG1 vOTUs were predicted through a combination of computational host prediction approaches via CRISPR-spacer and tRNA matching, according to Edwards *et al*. [35]. First, we constructed a reference host database (hereafter defined as RUGs), including rumen prokaryotic genomes from the Hungate1000 project [36] and rumen metagenome-assembled genomes (MAGs) from Stewart *et al.* [37], Anderson *et al.* [38], and Sato *et al.* [17]. RUGs were taxonomically annotated using GTDB-tk v.2.1.1 [39]. CRISPR-spacer sequences in RUGs were identified using CRISPRCasFinder v.4.2.20 [40]. CRISPR spacers with evidence levels of 3 or 4 were used for further analysis (n = 48 658). The CRISPR spacers of the RUGs were combined with those downloaded from CRISPRCasdb (https://crisprcas.i2bc.paris-saclay.fr/Home/Download; n = 353 377). The CRISPR-spacer dataset was queried against RVG1 vOTUs using blastn-short from BLAST+ v2.9.0 [41], allowing only one mismatch or gap with ≥95% identity of the spacer length. Identification of tRNA from RUGs and RVG1 vOTUs was performed using tRNAscan-SE 2.0 v.2.0.7 [42] with the general tRNA model, predicting 311 896 and 6 424 tRNAs, respectively. The retained tRNAs from the RUGs were merged with the prokaryotic tRNA database downloaded from GtRNAdb [43] (n = 260 923) and compared with tRNAs from RVG1 vOTUs using BLASTn [41]. Only a perfect match (100% identity and 100% length) was considered to imply host-virus interactions. To investigate whether RVG1 vOTUs were associated with hosts in multiple phyla, the following procedures were performed: (I) if more than two hosts of different phyla were predicted by the CRISPR approach, we decided that the RVG1 vOTUs may be associated with hosts in multiple phyla. (II) if different hosts at the phylum level were predicted between the two approaches, the result of the CRISPR method was adopted because the accuracy of the tRNA match was lower than that of the CRISPR-spacer match [35]. (III) if multiple phyla were predicted using tRNA methods, the phylum of the host was defined as “Undetermined.”

#### Read mapping and analysis of relative abundance and diversity

The cleaned reads of VLPs sequencing were mapped to RVG1 vOTUs using BamM “make” v1.7.3 (https://github.com/ecogenomics/BamM), and the reads that aligned over ≥80% of their length at ≥95% nucleic acid identity were obtained via the BamM “filter.” After filtering, contig depth coverage values were calculated using BamM “parse” with the “tpmean” algorithm. Contig breadth coverage was calculated using BEDtools “genomecov” v2.30.0 [44]. The abundance of contigs covered at <80% was considered zero. Finally, the retained contig depth coverage was normalized by the number of filtered reads in each sample. Normalized depth coverage was used to calculate the relative abundance of RVG1 vOTUs.

The accumulation curve showing the relationship between the number of RVG1 vOTUs and samples was calculated using the specaccum function with 1 000 permutations in the “vegan” package of R [45]. For alpha diversity analysis, Shannon diversity [46] and the number of RVG1 vOTU were estimated using the “vegan” package [45]. For beta diversity analysis, non-metric dimensional scaling analysis based on Bray–Curtis dissimilarity was performed. To evaluate the difference in the rumen virome between JB and F1, we compared the viral communities of the two breeds that were fed identical diets and kept in a single barn [16] using the Mann–Whitney U test and permutational multivariate analysis of variance (PERMANOVA) with 9 999 permutations for alpha and beta diversity, respectively, because the viral population in the rumen varied according to diet [7].

#### Metagenome sequencing

To further understand the viral-host relationship, we investigated the prokaryotic diversity of the samples using metagenomic sequencing. Raw sequenced data from 22 samples were downloaded from DDBJ (accession numbers: DRA011676 and DRA014084) and trimmed and filtered according to previous studies [16, 17]. Before mapping, RUGs were dereplicated at the species level [95% average nucleotide identity (ANI)] using dRep v.3.2.2 [47] to prevent arbitrary mapping between similar genomes [48]. The filtered reads were mapped against representative RUGs using BamM (https://github.com/Ecogenomics/BamM). The relative abundance was calculated using the CoverM v.0.4.0 “genome” (https://github.com/wwood/CoverM) with the following parameters: −m relative_abundance; -- min-read-percent-identity 0.95;--min-read-aligned-percent 0.75. We regarded the relative abundance of representative genomes as that of similar genomes (ANI > 95%).

#### Functional analysis of host genomes

Proteins of host prokaryotes were predicted using Prodigal v.2.6.3 [27]. The Carbohydrate-Active enZyme (CAZyme) annotation was performed using HMMER v.3.3 [32] against the dbCAN HMM v11 database with an E-value of < 1e-15 and coverage of > 0.35 [49].

## Results and discussion

### Overview of the rumen virome

As a result of VLP counting, the number of VLPs in the rumen was 1.6 ×10^10^ / mL, ranging from 2.0 ×10^9^ to 4.7 ×10^10^ / mL, comparable to a previous study [3, 4]. Sato *et al*. [16] reported that the 16S rRNA gene copy number of the total bacteria were 9.80 and 9.81 log_10_ copies/mL (6.3 ×10^9^ and 6.5 ×10^9^ copies/ml) in the JB and F1 cattle rumen, respectively. Based on these results, the number of viruses was the same or slightly higher than that of bacteria in the rumen, suggesting that prokaryotic cells in the rumen are constantly exposed to virus attacks, which may contribute to the regulation and control of the dynamics of the rumen microbiota.

The genome size of a virus is generally too small compared to that of its hosts, such as prokaryotes, resulting in a small percentage of viral DNA in the microbial community of the sample [50]. Therefore, the purification of VLPs is a crucial step before metagenomic sequencing. We used serial filtration and CsCl density gradient centrifugation for the rumen VLP purification, which were performed for samples in other environments, such as the human gut [51], the marine [52], and soil [53]. After the rumen VLPs sequencing, assembling, and dereplication, we retained 56 947 nonredundant contigs (≥ 5 kb). Among them, 52 293 contigs (91.8%) were classified as viral candidates via Virsortor2, suggesting that our laboratory procedure for VLP enrichment successfully removed cellular organism contamination and was more suitable for the metagenomic sequencing of VLPs in the rumen.

Subsequently, we examined the genome completeness of the vOTUs using CheckV, revealing that 8 232 vOTUs were predicted as ≥50% completeness, including 982 complete, 2 085 high-quality, and 5 165 medium-quality genomes (Supplementary Fig. 1). The length of RVG1 vOTUs ranged from 5 347 to 414 262 bp, with an average length of 39.5 ± 29.30 kb (mean ± SD) and GC content of 46.7 ± 8.43% (Supplementary data 1). The length of 42 RVG1 vOTUs were ≥ 200 kb, indicating that RVG1 vOTUs were possibly jumbo phages [54]. The 3 423 RVG1 vOTUs (41.6%) had genes encoding integrase, genes encoding recombinase, or both, suggesting they may be lysogenic phages (Supplementary Data 1). This result indicates that the rumen virome has lysogenic signatures and agrees with the human gut, where lysogenic phages dominate [51].

### Viral cluster and taxonomical assignment

To examine the relationship between RVG1 vOTUs and other viral genomes (RVG2 and RefSeq), a viral cluster analysis using vConTACT2 was conducted (Fig 1A; Supplementary data 2). In the network, RVG1 vOTUs were separated from the RefSeq virus genomes. The viral genomes in the entire dataset (RVG1, RVG2, and RefSeq) were classified into 1 146 VCs. Only 20 VCs were shared between RVG1 and RefSeq, whereas 260 VCs were shared between RVG1 and RVG2, suggesting rumen has a distinct virome from other environments. (Fig. 1B). The largest VC, VC_157, consisted of 271 viral genomes, with 251 RVG1 vOTUs. VC_157 included *Streptomyces* phage Bing (NC_042106) and *Streptomyces* phage *DrGrey* (NC_042016), which were isolated from a freshwater lake and dirt according to the Actinobacteriophage Database [55], respectively, suggesting that phages in VC_157 may be ubiquitous in the terrestrial biome. Most RVG1 vOTUs clustered with RVG2, not including RefSeq (3 047 vOTUs; 37.0%) or among RVG1 (2 116 vOTUs; 25.7%), and were singletons (2 539 vOTUs; 30.8%), indicating that most of the RVG1 vOTUs were rumen-specific phages. From these results, 4 655 RVG1 vOTUs were putative new genera that considerably expanded the rumen virus genomes. Furthermore, we performed a ViPTree analysis [29] against the RVG1 vOTUs that were not clustered with RefSeq viral genomes via vConTACT2 to assign taxonomy. Combined with the results from the two tools, a total of 1 053 RVG1 vOTUs (12.8%) were taxonomically assigned at the family level. Among them, Siphoviridae (710 vOTUs) was the most abundant, followed by Myoviridae (238 vOTUs) and Podoviridae (97 vOTUs), which were classified within the order Caudivirales (Fig. 1C). The 231 Siphoviridae vOTUs (32.5%) and 126 Myoviridae vOTUs (52.9%) were classified as lysogenic viruses. On the other hand, all Podoviridae vOTUs lacked lysogeny genes, in disagreement with the human fecal virome, where the family was composed of both lytic and lysogenic members, and dominant representatives were lysogenic viruses [51]. Viruses assigned to these three families have been found in the rumen of cattle [6, 7], moose [8], goats, and sheep [56], suggesting that these are the dominant families in the rumen virome. In contrast, most RVG1 vOTUs were classified as “unassigned,” consistent with previous studies on virome in other environments [13, 57] and suggest the necessity of further expansion of viral genomes, especially those of rumen viruses.

**Fig. 1.**
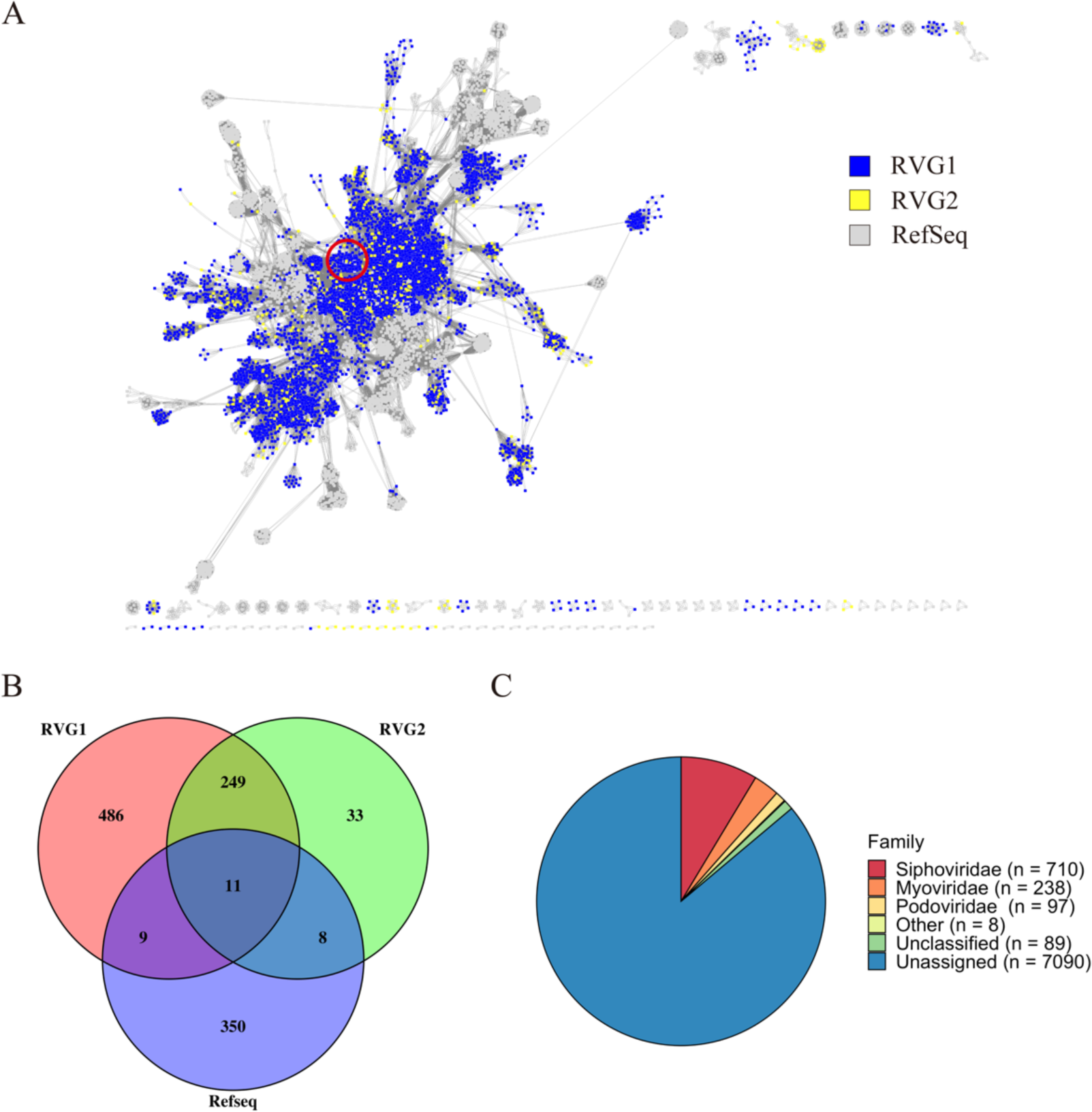
Interaction between RVG1 and other viral genomes, and taxonomy of RVG1. (A) Viral cluster analysis with vConTACT2 using a gene-sharing network. The RVG1 dots represent vOTUs reported by this study. The RVG2 dots represent vOTUs constructed from the rumen virome reported by Anderson *et al.* [7] and Solden *et al.* [8]. The red circle shows VC_157, which is the largest VC. (B) Venn diagram of shared viral clusters among RVG1, RVG2, and RefSeq. (C) Taxonomic diversity of the RVG1vOTUs. The pie chart shows the taxonomic classification of RVG1 vOTUs at a family level. “Unclassified” indicates the RVG1 vOTUs assigned to more than two families.

#### Virus-host interaction

We used two methods to identify potential virus-host interactions: CRISPR spacers and tRNA sequence match, which utilize nucleotide sequence similarity [35]. Overall, putative hosts were predicted for only 1 223 RVG1 vOTUs (14.9%) (Supplementary Data 3). Notably, putative hosts of 1 146 RVG1 vOTUs included at least one genome derived from RUGs, including 146 MAGs reconstructed from the rumens of the same animals used in this study [17]. These results strongly suggested that rumen viruses mainly infect rumen-specific bacteria and archaea. Some rumen phages (531 vOTUs) were related to multiple prokaryotic hosts, although most were associated with hosts in the same phylum (Fig. 2A; Supplementary Data 3). Additionally, some putative hosts in RUGs (1 088 prokaryotic genomes) were associated with only one RVG1 vOTU, whereas the others (1 179 prokaryotic genomes) were related to more than two RVG1 vOTUs (max: 33) (Fig. 2A; Supplementary data 3). As different viruses have different mechanisms of infection in hosts [58], it is difficult for hosts attacked by multiple viruses to acquire complete resistance to any virus. Thus, having multiple virus-host pairs may be crucial to prevent the host from developing resistance to specific viruses and to help regulate and control the prokaryotic population in the rumen. We expected that the putative hosts of most RVG1 vOTUs were the MAGs reported by Sato *et al*. [17] because they were reconstructed from the whole metagenome dataset in the rumen of the same animals used for VLPs sequencing in the present study; however, only 69 RVG1 vOTUs were related to the MAGs. One possible reason is that the number of MAGs (146) in the previous study was too small.

**Fig. 2.**
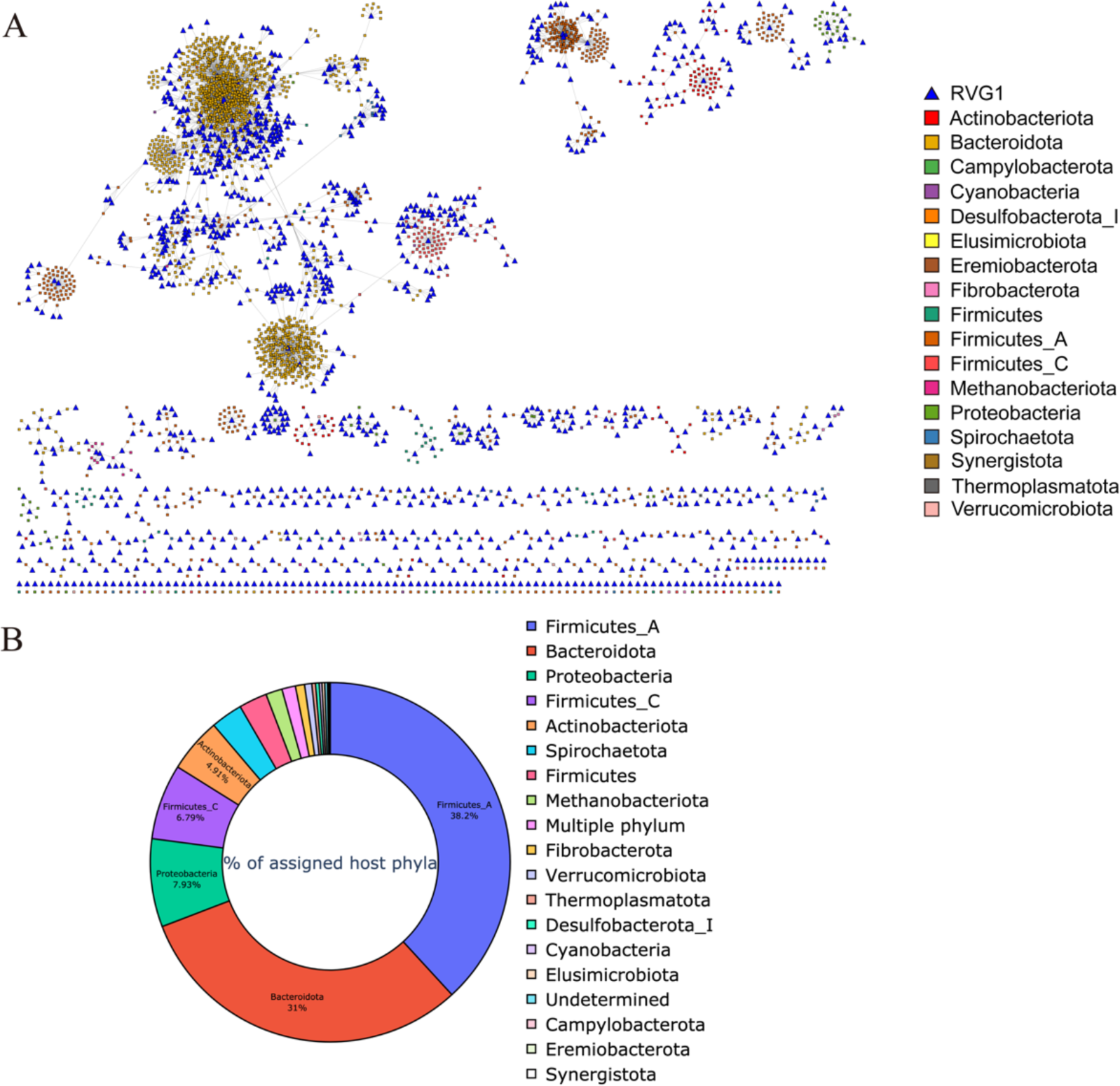
Virus-host interactions and taxonomy of the hosts. (A) Relationships of RVG1 vOTUs and the host. (B) Taxonomy of the rumen prokaryote genomes infected by RVG1 vOTUs. Taxonomy was assigned using GTDB-tk v.2.1.1 [39].

Most RVG1 vOTUs, whose hosts were predicted, had a narrow host range, consistent with the results for viruses in other environments [15, 57, 59]. Only 15 RVG1 vOTUs were potentially related to multiple phyla (Fig. 2B; Supplementary data 1). Among them, three vOTUs were associated with similar phyla in Firmicutes (Firmicutes, Firmicutes_A, and Firmicutes_C according to the GTDB), whereas the others were associated with multiple hosts in distant phyla. Considering that generalists with a broad host range spanning multiple phyla were found in the human gut when using the *in silico* approach [13, 60], RVG1 vOTUs may be able to infect hosts in multiple distinct phyla. However, it is necessary to conduct further *in vivo* experiments to investigate whether there are phages related to multiple phyla in the rumen. The predicted prokaryotic hosts spanned two archaeal and 15 bacterial phyla. Most bacterial hosts of RVG1 vOTUs belonged to Firmicutes (Firmicutes, Firmicutes_A, and Firmicutes_C according to GTDB; 47.5%) and Bacteroidetes (Bacteroidota according to GTDB; 31.0%), which are predominant phyla in the rumen [61, 62, 63, 64]. Notably, many vOTUs that were connected to Firmicutes_A and Bacteroidota were associated with the family Lachnospiraceae (222 vOTUs) and the genus *Prevotella* (190 vOTUs), respectively (Supplementary data 3). This might be associated with the results of Sato *et al*. [17], in which many Lachnospiraceae and *Prevotella* MAGs were obtained from rumen metagenomic data of the animals used in this study. *Prevotella* is the predominant genus in the rumen [65], and Lachnospiraceae is an essential family in the JB rumen [16, 66].

In this study, we identified 22 RVG1 vOTUs with archaeal hosts. The archaeal RVG1 vOTUs were grouped into three clades in the proteomic tree generated using ViPTree [29] with archaeal viruses in RefSeq (Supplementary Fig 2). Two clades (Clades 1 and 2) included 17 RVG1 vOTUs, of which the putative host was *Methanobrevibacter* (*Methanobrevibacter, Methanobrevibacter_A*, and *Methanobrevibacter_B* according to GTDB). Most RVG1 vOTUs in Clade 2 were clustered by vConTACT2 into VC_1409, VC_1443, and VC_1445, whereas the vOTUs in Clade 1 belonged to a variety of VCs, and the vOTUs did not have a lysogenic signature. In contrast, the other clade (Clade 3) consisted of four vOTUs related to *Methanomethylophilus* and clustered into VC_1245. In the rumen, *Methanobrevibacter* spp. is a dominant methanogen [67], whereas *Methanomethylophilus* spp. is a methylotrophic archaeon negatively correlated with methane emissions [68]. Therefore, the archaeal vOTUs in RVG1 probably control archaeal populations in the rumen and affect methane production in the rumen.

### Potential auxiliary metabolic genes

Viruses mediate the metabolism of hosts by AMGs. A total of 2 764 putative AMGs were identified from 1 720 RVG1 vOTUs, including 587 RVG vOTUs that have more than two AMG (Supplementary data 1). The AMG have been summarized in Supplementary Data 4. The AMGs included 1 026, 557, 403, 8, and 2 putative AMGs classified as “Miscellaneous (MISC),” “Carbon utilization,” “Organic nitrogen,” “Transporters,” and “Energy,” respectively (Fig. 3A). CAZymes are involved in the breakdown of complex carbohydrates and glycoconjugates [69], and a wide diversity of glycoside hydrolases (GHs) are involved in lignocellulose degradation in the rumen [70]. A variety of AMGs encoding GHs were found in 67 RVG1 vOTUs, including GH2, GH5, GH16, GH25, GH28, GH29, GH33, GH43, GH74, GH87, GH113, GH114, and GH141. Interestingly, the highest number of AMGs encoding the GH family was GH114 (endo-α-1,4-polygalactosaminidase) in 30 RVG1 vOTUs (Supplementary Data 4). Additionally, eight of nine RVG1 vOTUs belonging to VC_1309 harbored GH114 AMGs. Although GH114 has no or low abundance in the rumen [70, 71], and there is little knowledge of how GH114 contributes to feed digestion, it may have an important role. Fifteen bacterial hosts, including *Ruminococcus* and *Prevotella* predominantly responsible for fiber degradation in the rumen [72], interacted with RVG1 vOTUs that have GH AMGs. We investigated genes encoding GHs in putative host genomes to elucidate the ability of the host to degrade carbohydrates. A variety of GHs were found in all genomes, ranging from 15 to 136 GHs (Supplementary Fig. 3). According to these results, the RVG1 vOTUs that have AMGs encoding GHs may improve feed digestion in the rumen to augment the carbohydrate metabolism of fiber-degrading bacteria. In addition, 624, 254, and 248 AMGs encoded dUTP pyrophosphatase, DNA (cytosine-5)-methyltransferase 1, and ribonucleoside-triphosphate reductase, respectively (Fig. 3B).

**Fig. 3.**
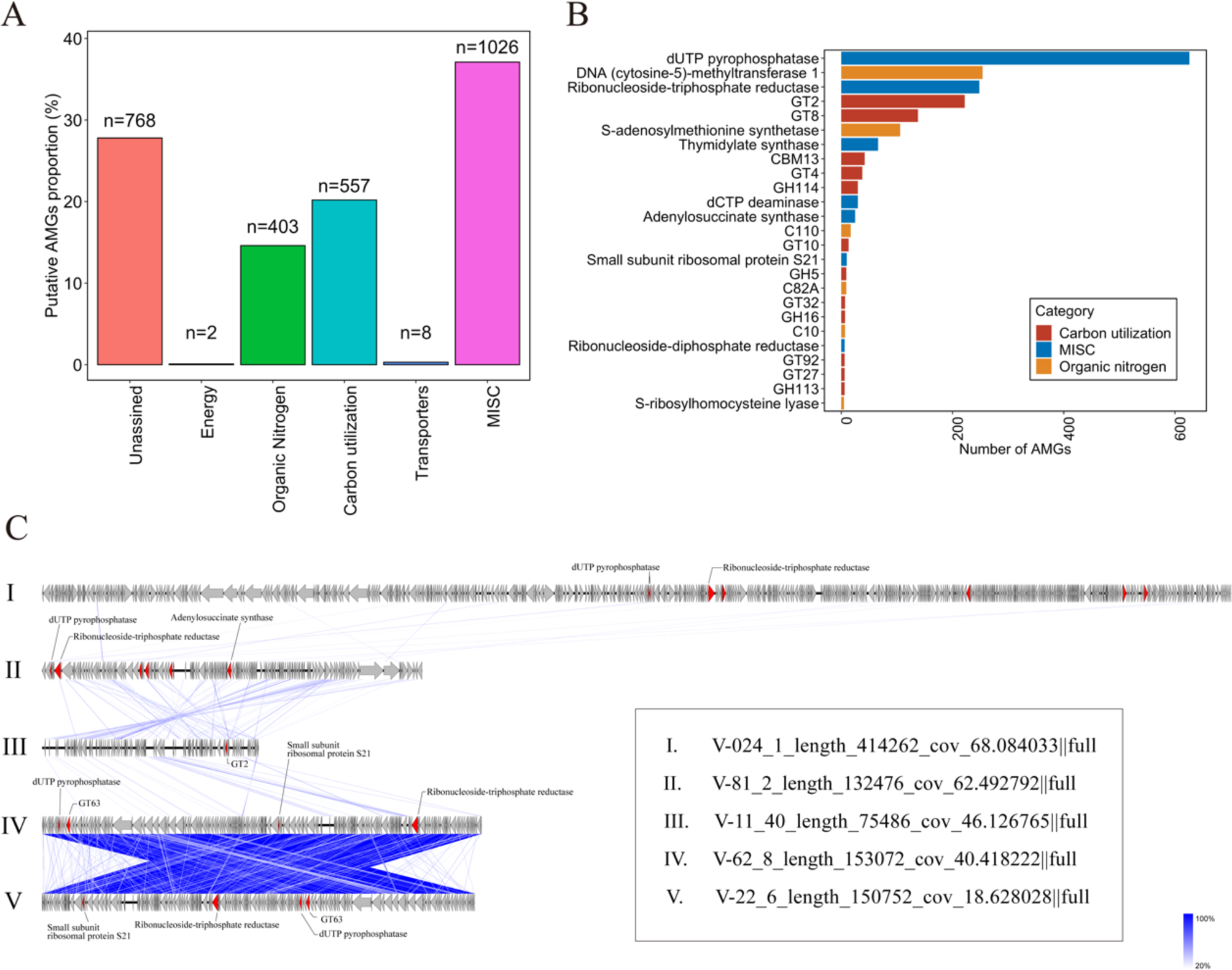
The auxiliary metabolic genes (AMGs) found in RVG1 vOTUs and genome map of archaeal RVG1 vOTU that have AMGs. (A) Bar plot of the number of AMGs in different categories. (B) The bar plot shows the number of AMGs found more than five times in RVG1 vOTUs. The AMGs were identified and categorized using DRAM-v [33]. (C) Linear genome map of archaeal RVG1 vOTUs in clade 1 that have AMGs. The red arrows show AMGs predicted with DRAM-v [33]. The blue regions between the two genomes indicate the level of tBLASTx identity.

Notably, 25 AMGs were found in eight RVG1 vOTUs that are associated with archaea, mostly *Methanobrevibacter* (*Methanobrevibacter* and *Methanobrevibacter_A* according to the GTDB) (Fig. 3C; Supplementary Fig. 4). Most archaeal RVG1 vOTUs in Clade 1 had more than two AMGs (Fig. 3C). Many AMGs in the archaeal RVG1 vOTUs in clade 1 encoded dUTP pyrophosphatase and ribonucleoside-triphosphate reductase (Fig. 3C). dUTP pyrophosphatase hydrolyzes dUTP to dUMP, preventing the misincorporation of uracil into DNA [73], whereas ribonucleoside-triphosphate reductase is involved in the catalysis of ribonucleotides into deoxyribonucleotides, which is essential for DNA synthesis and repair. Thus, archaeal RVG1 vOTUs likely accelerate DNA metabolism in hosts during infection.

### Individual rumen virome and viral diversity

After mapping VLP sequencing reads, the RVG1 vOTUs recruited 46.3 ± 6.30% reads per sample (mean ± SD). Moreover, the accumulation curve of RVG1 vOTUs did not plateau (Fig. 4A). These results indicate that RVG1 vOTUs did not completely cover rumen viral diversity. Notably, 4 212 (51.2%) (defined as “individual-specific” vOTUs) and 2 020 RVG1 vOTUs (24.5%) were present in one and two samples, respectively, while only 42 vOTUs (defined as “core” vOTUs) were found in more than 50% samples (Fig. 4B). This implies that rumen viruses are highly individual-specific, consistent with human gut viruses [74, 75]. The relative abundance of “individual-specific” vOTUs was 23.8 ± 5.08%, while “core” vOTUs was only 8.6 ± 1.84% (mean ± SE) (Fig. 4C). This suggests that individual-specific vOTUs are more important than “core” vOTUs in the rumen virome. In the rumen virome, Siphoviridae (8.3 ± 0.64%) was predominant, followed by Myoviridae (2.5 ± 0.34%) and Podoviridae (2.2 ± 0.90%) (mean ± SE) (Fig. 4D). Additionally, the relative abundance of lysogenic vOTUs was 34.4 ± 1.63% (mean ± SE), indicating that the abundance of many hosts, such as bacteria and archaea, are regulated via lysogeny in the rumen.

**Fig. 4.**
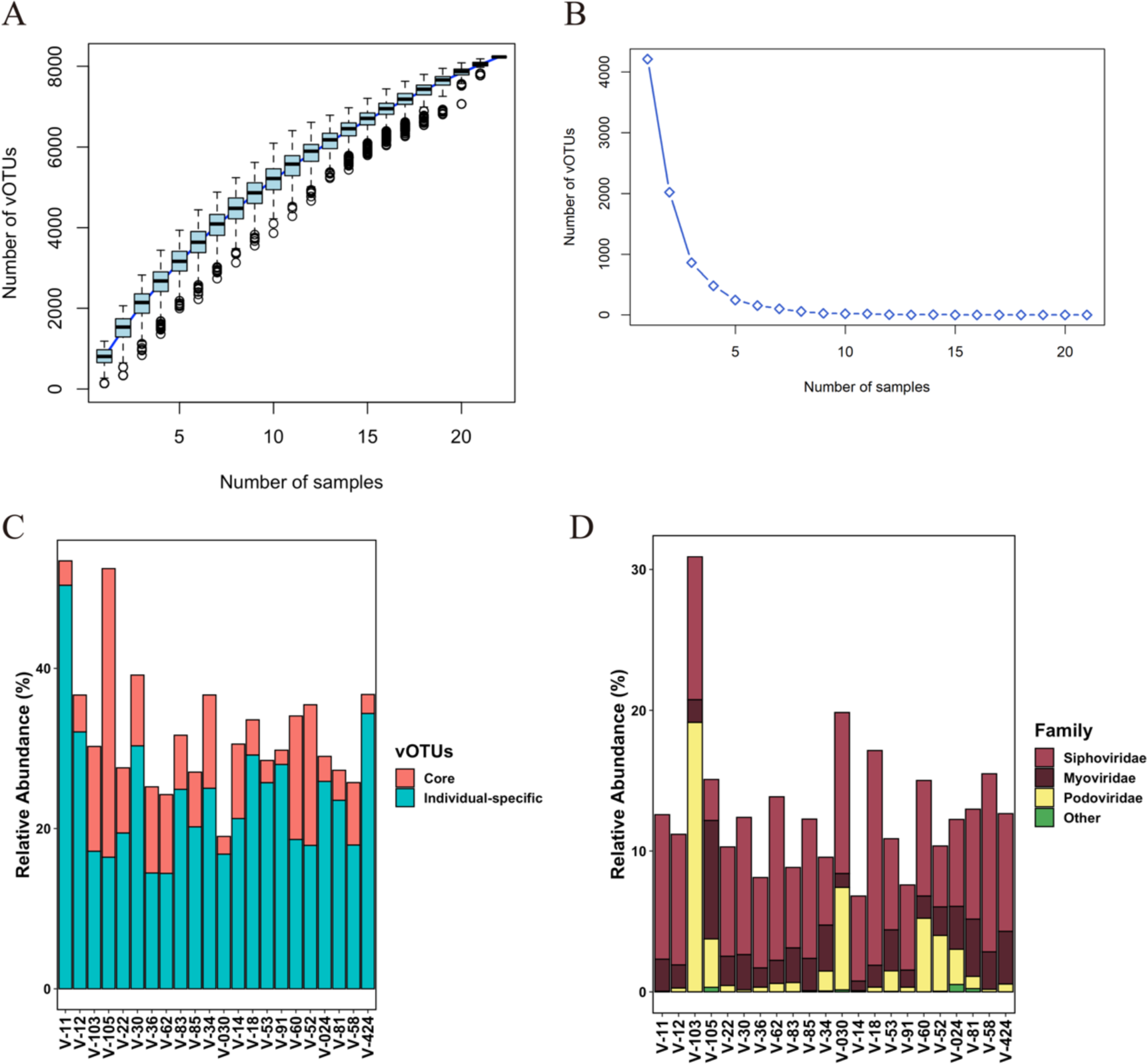
Characteristics of individual rumen virome. (A) Accumulation curve of RVG1 vOTUs. (B) Curve showing the number of RVG1 vOTUs that were present in the number of rumen samples. (C) Relative abundance of “core” and “individual” vOTUs in samples. (D) Relative abundance of rumen viromes at the family level.

Next, we investigated the degree of virus-host co-occurrence of RVG1 vOTUs with putative hosts in the RUGs (Supplementary Fig. 5). As a result, 428 vOTUs and their putative hosts simultaneously existed (Group 1), whereas 718 vOTUs were not present in their hosts in the identical environments (Group 2). This suggests that some rumen viruses interact with prokaryotes that do not include the RUGs. Considering that the putative hosts of most RVG1 vOTUs were not estimated, reconstructing a higher number of rumen MAGs is needed for future studies on virus-host interactions in the rumen.

In the human gut, many factors (e.g., age, diet, and health) influence virome composition [12, 76]. Regarding the rumen virome, however, diet is the only factor affecting the rumen virome uncovered by the previous study thus far [7]. Therefore, examining whether other factors influence the rumen virome is necessary to expand our knowledge. To investigate the effect of cattle breed on rumen viral diversity, we compared the rumen virome between JB and F1 steers fed identical diets and kept in a single cattle barn [16] by examining α-and β-diversity. Interestingly, the rumen viral community structure (β-diversity) was significantly different between JB and F1 steers (P = 0.003), although no difference was found in α-diversity (P > 0.05) (Fig. 5; Supplementary Fig. 6). This agrees with a previous study that discovered differences in prokaryotic community structures between the JB and F1 rumen [16]. This indicates that the cattle breed, as with diet, is a factor that influences the rumen virome and the rumen prokaryotic community.

**Fig. 5.**
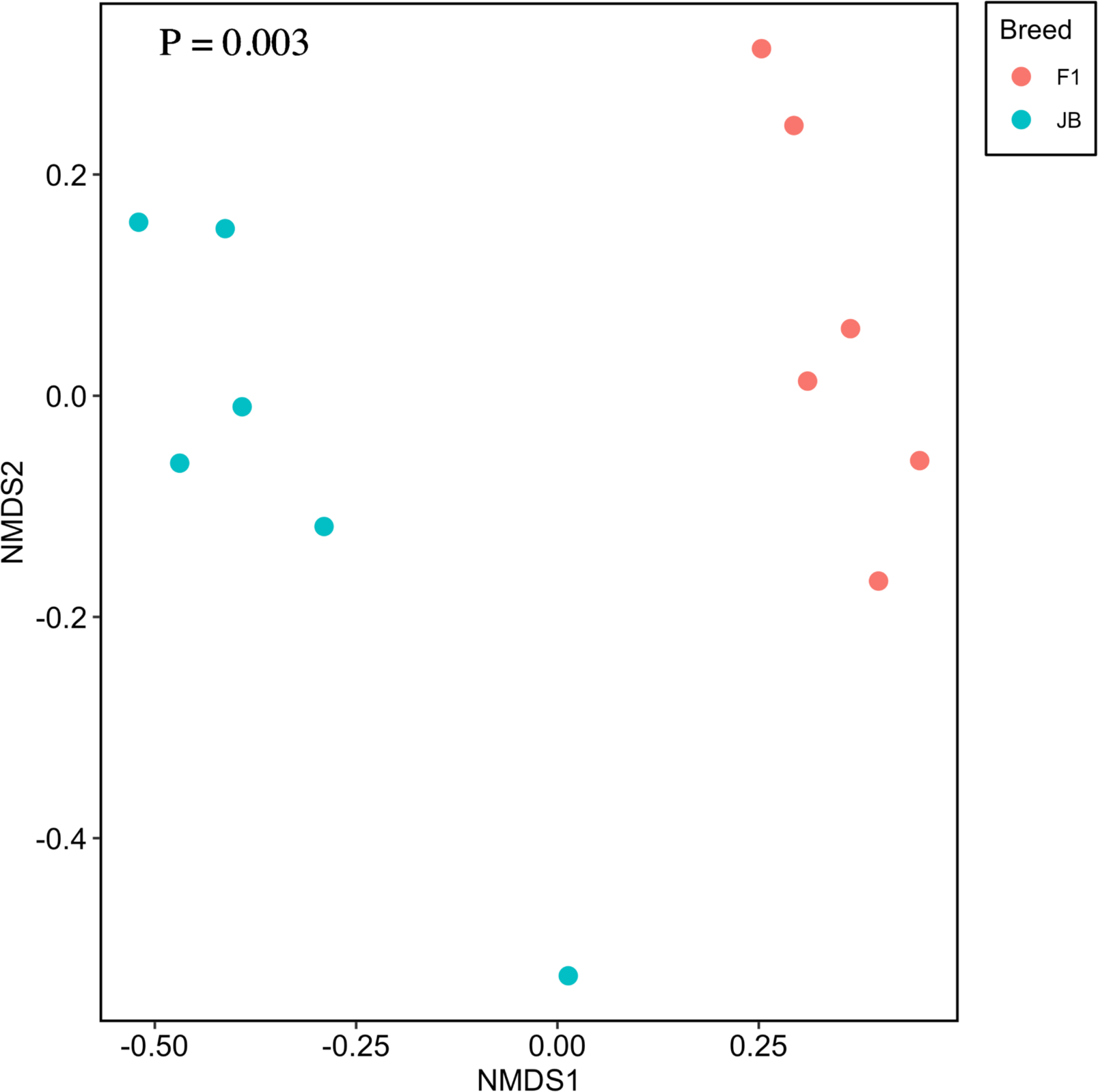
Non-metric multidimensional scaling plots based on Bray–Curtis dissimilarities in RVG1 vOTUs. The Japanese Black (JB) and Japanese Black × Holstein crossbred (F1) steers were fed identical diets and kept in a single cattle barn [16].

## Conclusion

In summary, we constructed the first genome catalog of the rumen virus of cattle, including 8 232 vOTUs that were classified as “Complete,” “High-quality,” and “Medium-quality” by CheckV. These rumen viral genomes are significantly helpful for future studies on rumen viromes using metagenomic sequencing.

We discovered that most rumen viruses were highly rumen-and individual-specific and were related to rumen-specific prokaryotes. In addition, the study revealed that the cattle breed influenced the rumen virome. These findings advance our understanding of rumen viruses. Furthermore, 22 RVG1 vOTUs associated with archaea were identified in the rumen. Methane is produced during microbial fermentation in the rumen by archaea, commonly known as methanogens, and contributes to global warming, leading to significant interest in developing methane reduction strategies. The phage therapy-based approach is considered one of the promising approaches for inhibiting enteric methane production from the rumen [77, 78, 79]. In fact, the bionanoparticle fused with the lytic enzyme (PeiR), which was found within the provirus genome of *Methanobrevibacter ruminantium* M1 [76], inhibited methane production in a continuous-flow rumen fermentation system [80]. To further develop this approach, understanding archaeal viruses is crucial. The genomic information of the archaeal vOTUs in this study helps to develop a feasible strategy for decreasing methane production from the rumen.

## Acknowledgements

This study was partly supported by grants from Kieikai Research Foundation to YS and JSPS KAKENHI Grant Number JP21H05057 to TY. The super-computing resource was provided by Human Genome Center, the Institute of Medical Science, the University of Tokyo.

## Data availability

Raw sequencing data of VLPs were deposited in DBBJ (accession number: DRA015871). The RVG1 vOTU and the other vOTUs which classified as “Low-quality” and “Undetermined” by CheckV are available at figshare (https://doi.org/10.6084/m9.figshare.22256155).

---

**Supplementary Fig. 1.**
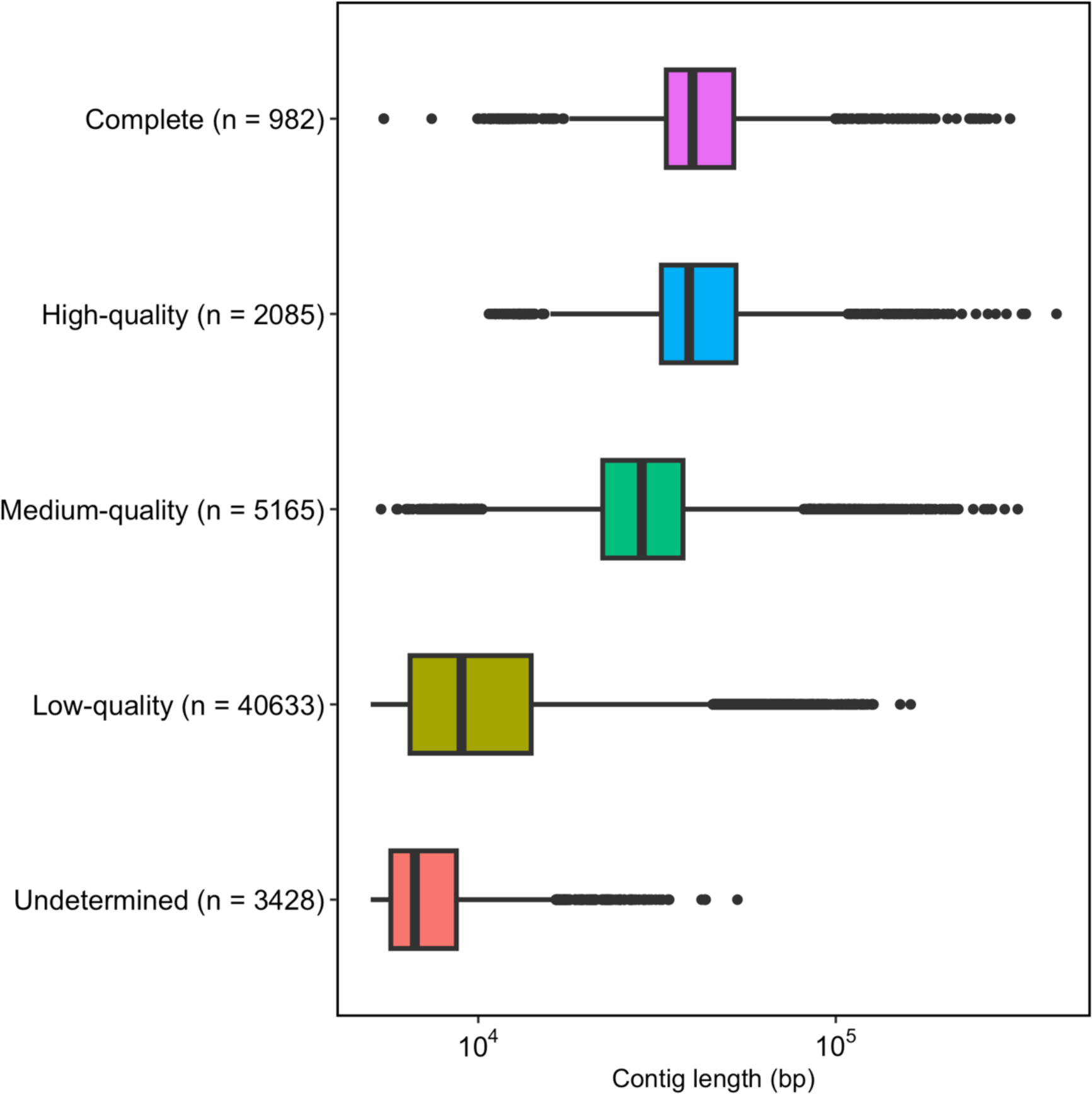
Distribution of contig length and quality tiers of the rumen viruses. The quality of the rumen virus genomes was assessed using CheckV [25]. “High-quality,” “Medium-quality,” and “Low-quality” indicate ≥90%, 50–90%, and 0–50% completeness, respectively.

**Supplementary Fig. 2.**
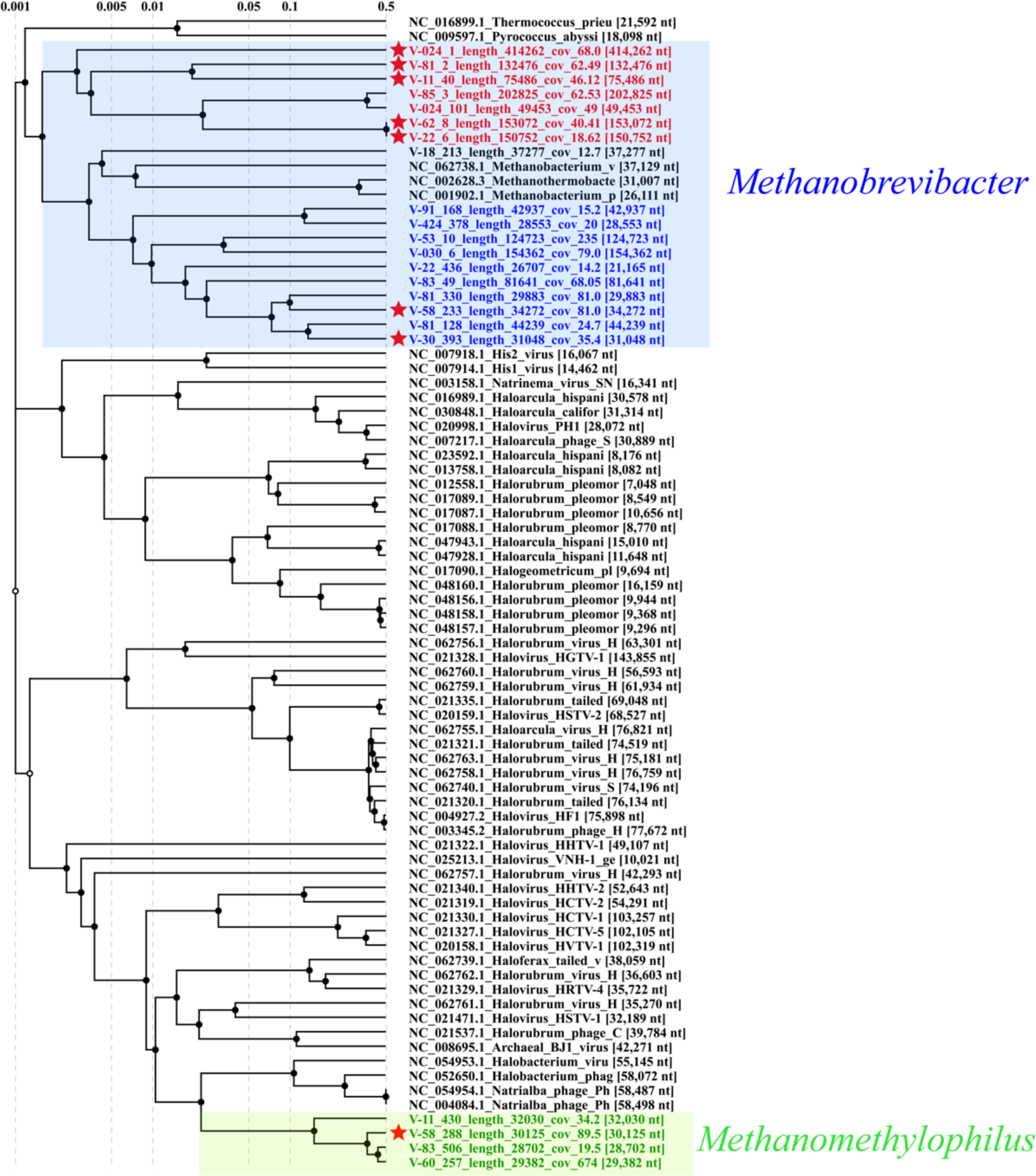
Proteomic tree with archaeal virus in RVG1 and RefSeq. Archaeal RVG1 vOTUs were roughly clustered in three clades (Red, Clade 1; Blue, Clade 2; Green, Clade 3). Clades 1 and 2 were related to *Methanobrevibacter* (*Methanobrevibacter*, *Methanobrevibacter_A*, and *Methanobrevibacter_B* according to GTDB), while Clade 3 was associated with *Methanomethylophilus.* The proteomic tree was generated by ViPTree [29]. The stars show the archaeal RVG1 vOTUs that have some AMGs.

**Supplementary Fig. 3.**
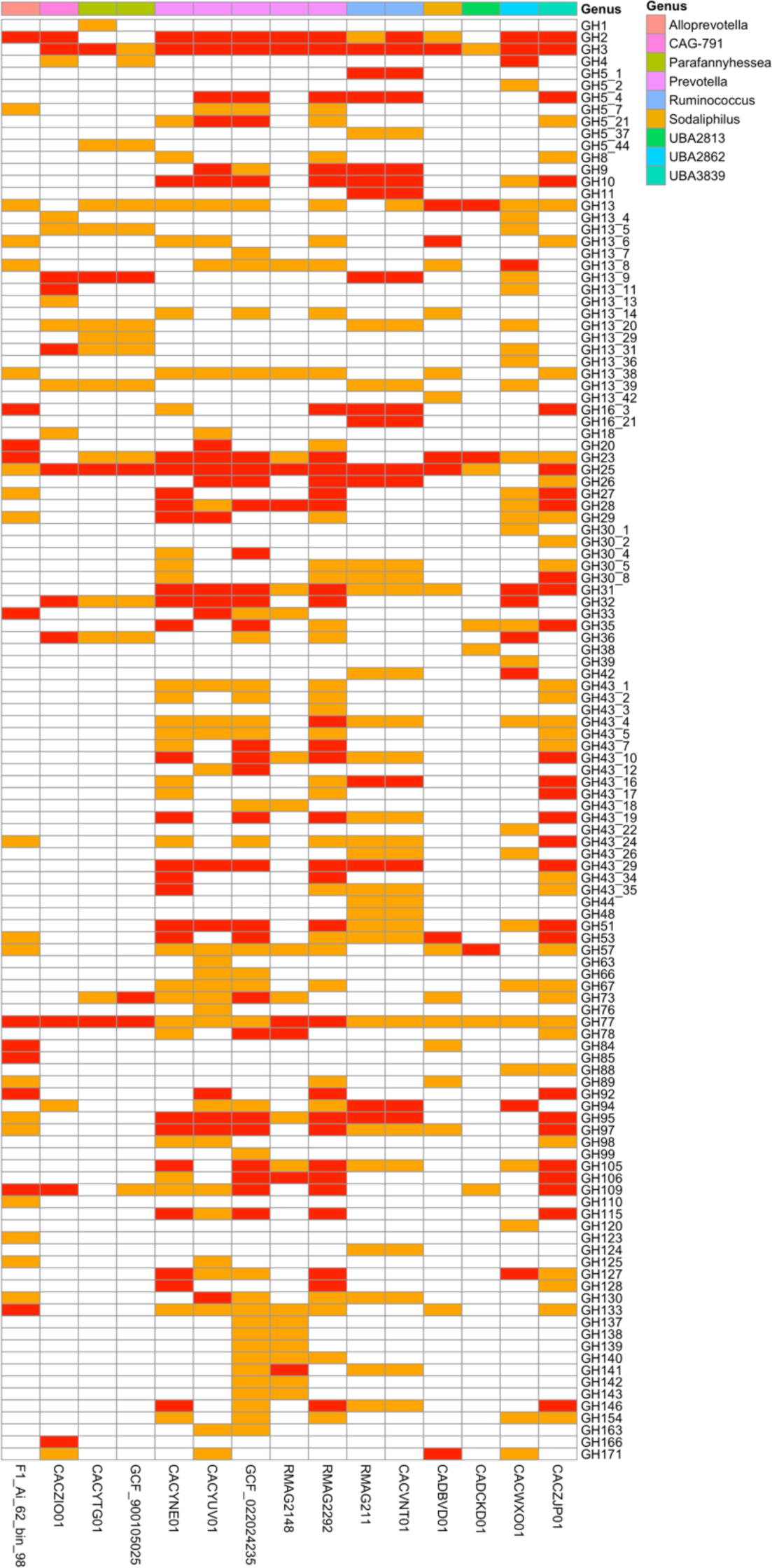
Heat map of genes encoding glycoside hydrolases (GH) in procaryotic genomes related to RVG1 vOTUs that have AMGs encoding GHs. The presence of more than two or only one GH gene is depicted in red and orange, respectively.

**Supplementary Fig. 4.**
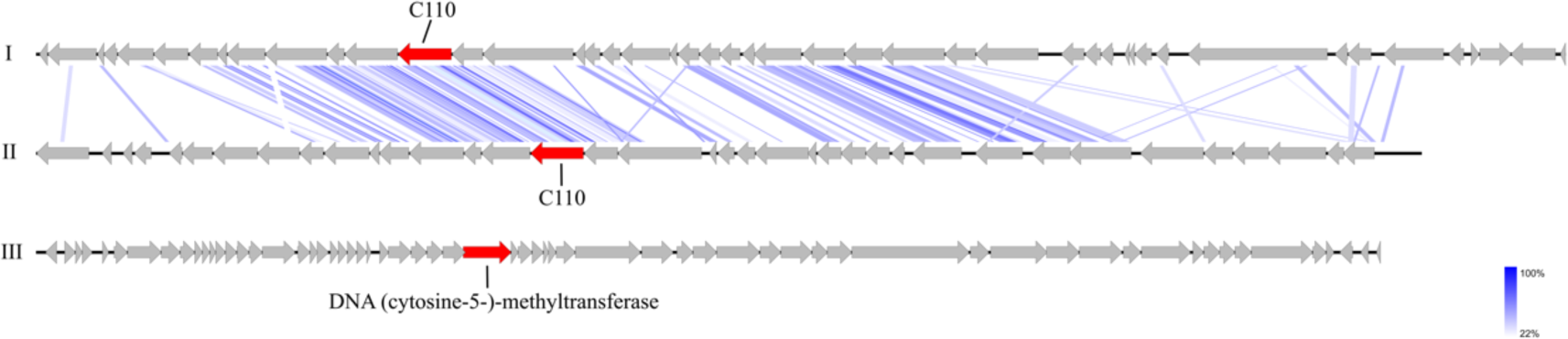
Linear genome map of archaeal RVG1 vOTUs in clades 2 and 3 that have AMGs. Genome map of I and II are V-58_233_length_34272_cov_81.087208||full and V-30_393_length_31048_cov_35.404478||full, respectively, in clade 2, and that of III is V-58_288_length_30125_cov_89.538211||full in clade 3. The red arrows show AMGs predicted with DRAM-v [33]. The blue regions between the two genomes indicate the level of tBLASTx identity.

**Supplementary Fig. 5.**
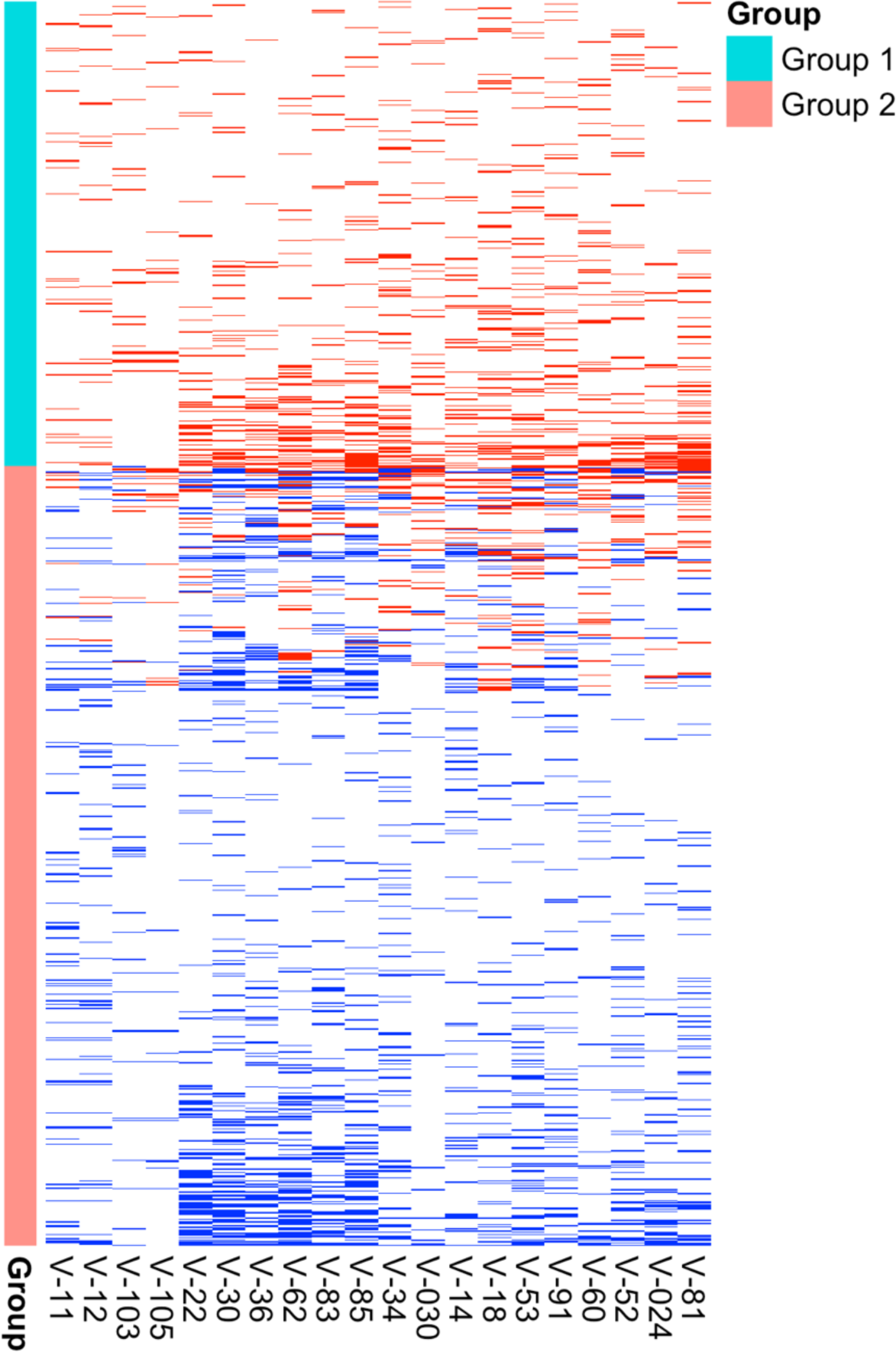
Heatmap showing the co-occurrence between RVG1 vOTUs and the host in the samples. Group 1 shows that RVG1 vOTUs are simultaneously present with their putative hosts in all samples (n = 428), while Group 2 indicates that RVG1 vOTUs and their hosts do not exist at least in one sample (n = 728). The red lines show the co-occurrence of RVG1 vOTUs and their hosts. The blue lines indicate that RVG1 vOTUs are present, but their hosts do not exist. Each row represents one RVG1 vOTUs, and each column represents one sample.

**Supplementary Fig. 6.**
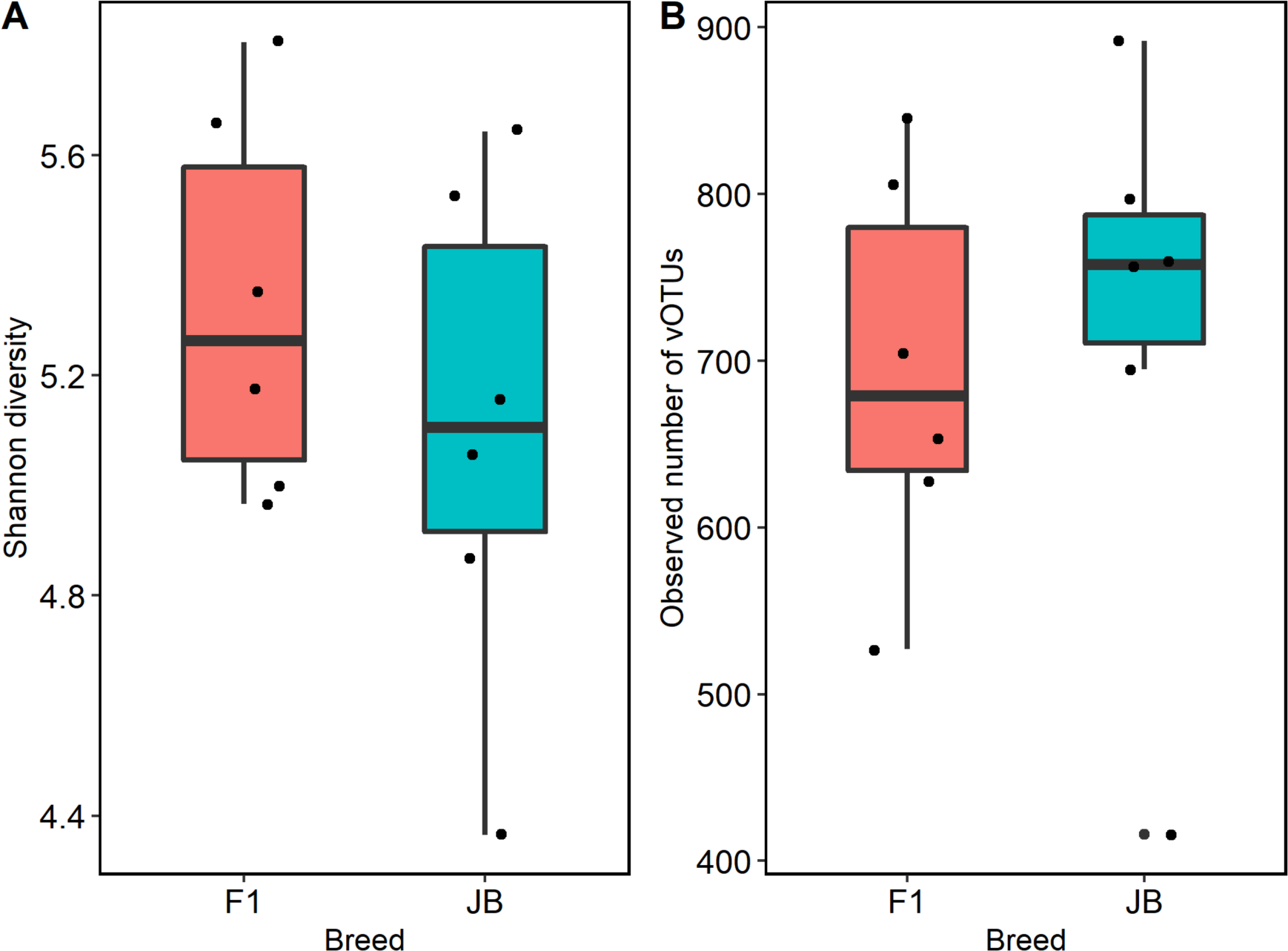
Alpha diversity in the rumen virome of Japanese Black (JB) and Japanese Black × Holstein crossbred (F1) steers. (A) Boxplot that shows Shannon diversity indices. (B) Observed number of RVG1 vOTUs. The JB and F1 steers were fed identical diets and kept in a single cattle barn [16].

## REFERENCES

[1] Morgavi DP, Kelly WJ, Janssen PH, Attwood GT. Rumen microbial (meta)genomics and its application to ruminant production. Animal 2013; 7:184–201.

[2] Huws SA, Creevey CJ, Oyama LB, Mizrahi I, Denman SE, Popova M, et al. Addressing global ruminant agricultural challenges through understanding the rumen microbiome: Past, present, and future. Front Microbiol 2018; 9:2161.

[3] Paynter MJB, Ewert DL, Chalupa W. Some morphological types of bacteriophages in bovine rumen contents. Appl Microbiol 1969; 18:942–943.

[4] Klieve A V., Swain RA. Estimation of ruminal bacteriophage numbers by pulsed-field gel electrophoresis and laser densitometry. Appl Environ Microbiol 1993; 59:2299–2303.

[5] Gilbert RA, Townsend EM, Crew KS, Hitch TCA, Friedersdorff JCA, Creevey CJ, et al. Rumen virus populations: technological advances enhancing current understanding. Front Microbiol 2020; 11:450.

[6] Berg Miller ME, Yeoman CJ, Chia N, Tringe SG, Angly FE, Edwards RA, et al. Phage–bacteria relationships and CRISPR elements revealed by a metagenomic survey of the rumen microbiome. Environ Microbiol 2012; 14:207–227.

[7] Anderson CL, Sullivan MB, Fernando SC. Dietary energy drives the dynamic response of bovine rumen viral communities. Microbiome 2017; 5:155.

[8] Solden LM, Naas AE, Roux S, Daly RA, Collins WB, Nicora CD, et al. Interspecies cross-feeding orchestrates carbon degradation in the rumen ecosystem. Nat Microbiol 2018; 3:1274–1284.

[9] Islam MM, Fernando SC, Saha R. Metabolic modeling elucidates the transactions in the rumen microbiome and the shifts upon virome interactions. Front Microbiol 2019; 10:2412.

[10] Ross EM, Petrovski S, Moate PJ, Hayes BJ. Metagenomics of rumen bacteriophage from thirteen lactating dairy cattle. BMC Microbiol 2013; 13:242.

[11] Roux S, Páez-Espino D, Chen IMA, Palaniappan K, Ratner A, Chu K, et al. IMG/VR v3: an integrated ecological and evolutionary framework for interrogating genomes of uncultivated viruses. Nucleic Acids Res 2021; 49:D764–D775.

[12] Gregory AC, Zablocki O, Zayed AA, Howell A, Bolduc B, Sullivan MB. The gut virome database reveals age-dependent patterns of virome diversity in the human gut. Cell Host Microbe 2020; 28:724–740.

[13] Camarillo-Guerrero LF, Almeida A, Rangel-Pineros G, Finn RD, Lawley TD. Massive expansion of human gut bacteriophage diversity. Cell 2021; 184:1098–1109.

[14] Nayfach S, Páez-Espino D, Call L, Low SJ, Sberro H, Ivanova NN, et al. Metagenomic compendium of 189,680 DNA viruses from the human gut microbiome. Nat Microbiol 2021; 6:960–970.

[15] Paez-Espino D, Eloe-Fadrosh EA, Pavlopoulos GA, Thomas AD, Huntemann M, Mikhailova N, et al. Uncovering Earth’s virome. Nature 2016; 536:425–430.

[16] Sato Y, Takebe H, Tominaga K, Oishi K, Kumagai H, Yoshida T, et al. Taxonomic and functional characterization of the rumen microbiome of Japanese Black cattle revealed by 16S rRNA gene amplicon and metagenome shotgun sequencing. FEMS Microbiol Ecol 2021; 97: fiab152.

[17] Sato Y, Takebe H, Oishi K, Yasuda J, Kumagai H, Hirooka H, et al. Identification of 146 metagenome-assembled genomes from the rumen microbiome of cattle in Japan. Microbes Environ 2022; 37:ME22039.

[18] Hurwitz BL, Deng L, Poulos BT, Sullivan MB. Evaluation of methods to concentrate and purify ocean virus communities through comparative, replicated metagenomics. Environ Microbiol 2013; 15: 1428–1440.

[19] Tillett D, Neilan BA. Xanthogenate nucleic acid isolation from cultured and environmental cyanobacteria. J Phycol 2000; 36: 251–258.

[20] Bolger AM, Lohse M, Usadel B. Trimmomatic: a flexible trimmer for Illumina sequence data. Bioinformatics 2014; 30: 2114–2120.

[21] Bankevich A, Nurk S, Antipov D, Gurevich AA, Dvorkin M, Kulikov AS, et al. SPAdes: a new genome assembly algorithm and its applications to single-cell sequencing. J Comput Biol 2012; 19: 455–477.

[22] Nurk S, Meleshko D, Korobeynikov A, Pevzner PA. MetaSPAdes: a new versatile metagenomic assembler. Genome Res 2017; 27: 824–834.

[23] Fu L, Niu B, Zhu Z, Wu S, Li W. CD-HIT: accelerated for clustering the next-generation sequencing data. Bioinformatics 2012; 28: 3150–3152.

[24] Guo J, Bolduc B, Zayed AA, Varsani A, Dominguez-Huerta G, Delmont TO, et al. VirSorter2: a multi-classifier, expert-guided approach to detect diverse DNA and RNA viruses. Microbiome 2021; 9: 37.

[25] Nayfach S, Camargo AP, Schulz F, Eloe-Fadrosh E, Roux S, Kyrpides NC. CheckV assesses the quality and completeness of metagenome-assembled viral genomes. Nat Biotechnol 2021; 39: 578–585.

[26] Bin Jang H, Bolduc B, Zablocki O, Kuhn JH, Roux S, Adriaenssens EM, et al. Taxonomic assignment of uncultivated prokaryotic virus genomes is enabled by gene-sharing networks. Nat Biotechnol 2019; 37: 632–639.

[27] Hyatt D, Chen G-L, LoCascio PF, Land ML, Larimer FW, Hauser LJ. Prodigal: prokaryotic gene recognition and translation initiation site identification. BMC Bioinformatics 2010; 11: 119.

[28] Shannon P, Markiel A, Ozier O, Baliga NS, Wang JT, Ramage D, et al. Cytoscape: a software environment for integrated models of biomolecular interaction networks. Genome Res 2003; 13: 2498–2504.

[29] Nishimura Y, Yoshida T, Kuronishi M, Uehara H, Ogata H, Goto S. ViPTree: the viral proteomic tree server. Bioinformatics 2017; 33: 2379–2380.

[30] Mihara T, Nishimura Y, Shimizu Y, Nishiyama H, Yoshikawa G, Uehara H, et al. Linking virus genomes with host taxonomy. Viruses 2016; 8: 66.

[31] Finn RD, Coggill P, Eberhardt RY, Eddy SR, Mistry J, Mitchell AL, et al. The Pfam protein families database: towards a more sustainable future. Nucleic Acids Res 2016; 44: D279–D285.

[32] Mistry J, Finn RD, Eddy SR, Bateman A, Punta M. Challenges in homology search: HMMER3 and convergent evolution of coiled-coil regions. Nucleic Acids Res 2013; 41: e121–e121.

[33] Shaffer M, Borton MA, McGivern BB, Zayed AA, La Rosa SL, Solden LM, et al. DRAM for distilling microbial metabolism to automate the curation of microbiome function. Nucleic Acids Res 2020; 48: 8883–8900.

[34] Sullivan MJ, Petty NK, Beatson SA. Easyfig: a genome comparison visualizer. Bioinformatics 2011; 27: 1009–1010.

[35] Edwards RA, McNair K, Faust K, Raes J, Dutilh BE. Computational approaches to predict bacteriophage–host relationships. FEMS Microbiol Rev 2016; 40: 258–272.

[36] Seshadri R, Leahy SC, Attwood GT, Teh KH, Lambie SC, Cookson AL, et al. Cultivation and sequencing of rumen microbiome members from the Hungate1000 Collection. Nat Biotechnol 2018; 36: 359–367.

[37] Stewart RD, Auffret MD, Warr A, Walker AW, Roehe R, Watson M. Compendium of 4,941 rumen metagenome-assembled genomes for rumen microbiome biology and enzyme discovery. Nat Biotechnol 2019; 37: 953–961.

[38] Anderson CL, Fernando SC. Insights into rumen microbial biosynthetic gene cluster diversity through genome-resolved metagenomics. Commun Biol 2021; 4: 818.

[39] Chaumeil P-A, Mussig AJ, Hugenholtz P, Parks DH. GTDB-Tk v2: memory friendly classification with the genome taxonomy database. Bioinformatics 2022; 38: 5315–5316.

[40] Couvin D, Bernheim A, Toffano-Nioche C, Touchon M, Michalik J, Néron B, et al. CRISPRCasFinder, an update of CRISRFinder, includes a portable version, enhanced performance and integrates search for Cas proteins. Nucleic Acids Res 2018; 46: W246–W251.

[41] Camacho C, Coulouris G, Avagyan V, Ma N, Papadopoulos J, Bealer K, et al. BLAST+: architecture and applications. BMC Bioinformatics 2009; 10: 421.

[42] Chan PP, Lin BY, Mak AJ, Lowe TM. tRNAscan-SE 2.0: improved detection and functional classification of transfer RNA genes. Nucleic Acids Res 2021; 49: 9077–9096.

[43] Chan PP, Lowe TM. GtRNAdb 2.0: an expanded database of transfer RNA genes identified in complete and draft genomes. Nucleic Acids Res 2016; 44: D184–D189.

[44] Quinlan AR, Hall IM. BEDTools: a flexible suite of utilities for comparing genomic features. Bioinformatics 2010; 26: 841–842.

[45] Oksanen J, Blanchet FG, Friendly M, Kindt R, Legendre P, McGlinn D, et al. Vegan: community ecology package (version 2.5-6). Compr R Arch Netw 2019.

[46] Shannon CE. A mathematical theory of communication. Bell Syst Tech J 1948; 27: 379–423.

[47] Olm MR, Brown CT, Brooks B, Banfield JF. dRep: a tool for fast and accurate genomic comparisons that enables improved genome recovery from metagenomes through de-replication. ISME J 2017; 11: 2864–2868.

[48] Robbins SJ, Song W, Engelberts JP, Glasl B, Slaby BM, Boyd J, et al. A genomic view of the microbiome of coral reef demosponges. ISME J 2021; 15: 1641–1654.

[49] Zhang H, Yohe T, Huang L, Entwistle S, Wu P, Yang Z, et al. dbCAN2: a meta server for automated carbohydrate-active enzyme annotation. Nucleic Acids Res 2018; 46: W95–W101.

[50] Reyes A, Semenkovich NP, Whiteson K, Rohwer F, Gordon JI. Going viral: next-generation sequencing applied to phage populations in the human gut. Nat Rev Microbiol 2012; 10: 607–617.

[51] Reyes A, Haynes M, Hanson N, Angly FE, Heath AC, Rohwer F, et al. Viruses in the faecal microbiota of monozygotic twins and their mothers. Nature 2010; 466: 334–338.

[52] Hurwitz BL, Sullivan MB. The Pacific Ocean Virome (POV): a marine viral metagenomic dataset and associated protein clusters for quantitative viral ecology. PLoS One 2013; 8: e57355.

[53] Trubl G, Jang H Bin, Roux S, Emerson JB, Solonenko N, Vik DR, et al. Soil viruses are underexplored players in ecosystem carbon processing. mSystems 2018; 3: e00076–18.

[54] Al-Shayeb B, Sachdeva R, Chen L-X, Ward F, Munk P, Devoto A, et al. Clades of huge phages from across Earth’s ecosystems. Nature 2020; 578: 425–431.

[55] Russell DA, Hatfull GF. PhagesDB: the actinobacteriophage database. Bioinformatics 2017; 33: 784–786.

[56] Namonyo S, Wagacha M, Maina S, Wambua L, Agaba M. A metagenomic study of the rumen virome in domestic caprids. Arch Virol 2018; 163: 3415–3419.

[57] Li Z, Pan D, Wei G, Pi W, Zhang C, Wang J-H, et al. Deep sea sediments associated with cold seeps are a subsurface reservoir of viral diversity. ISME J 2021; 15: 2366–2378.

[58] Hasan M, Ahn J. Evolutionary dynamics between phages and bacteria as a possible approach for designing effective phage therapies against antibiotic-resistant bacteria. Antibiotics 2022; 11: 915.

[59] Gu C, Liang Y, Li J, Shao H, Jiang Y, Zhou X, et al. Saline lakes on the Qinghai-Tibet Plateau harbor unique viral assemblages mediating microbial environmental adaption. Iscience 2021; 24: 103439.

[60] de Jonge PA, Wortelboer K, Scheithauer TPM, van den Born B-JH, Zwinderman AH, Nobrega FL, et al. Gut virome profiling identifies a widespread bacteriophage family associated with metabolic syndrome. Nat Commun 2022; 13: 3594.

[61] Kim M, Morrison M, Yu Z. Status of the phylogenetic diversity census of ruminal microbiomes. FEMS Microbiol Ecol 2011; 76: 49–63.

[62] Jami E, White BA, Mizrahi I. Potential role of the bovine rumen microbiome in modulating milk composition and feed efficiency. PLoS One 2014; 9: e85423.

[63] Pitta DW, Indugu N, Kumar S, Vecchiarelli B, Sinha R, Baker LD, et al. Metagenomic assessment of the functional potential of the rumen microbiome in Holstein dairy cows. Anaerobe 2016; 38: 50–60.

[64] Wang L, Xu Q, Kong F, Yang Y, Wu D, Mishra S, et al. Exploring the goat rumen microbiome from seven days to two years. PLoS One 2016; 11: e0154354.

[65] Henderson G, Cox F, Ganesh S, Jonker A, Young W, Abecia L, et al. Rumen microbial community composition varies with diet and host, but a core microbiome is found across a wide geographical range. Sci Rep 2015; 5: 14567.

[66] Ogata T, Makino H, Ishizuka N, Iwamoto E, Masaki T, Ikuta K, et al. Long-term high-grain diet altered the ruminal pH, fermentation, and composition and functions of the rumen bacterial community, leading to enhanced lactic acid production in Japanese Black beef cattle during fattening. PLoS One 2019; 14: e0225448.

[67] Janssen PH, Kirs M. Structure of the archaeal community of the rumen. Appl Environ Microbiol 2008; 74: 3619–3625.

[68] Martínez-Álvaro M, Auffret MD, Stewart RD, Dewhurst RJ, Duthie C-A, Rooke JA, et al. Identification of complex rumen microbiome interaction within diverse functional niches as mechanisms affecting the variation of methane emissions in bovine. Front Microbiol 2020; 11: 659.

[69] Cantarel BL, Coutinho PM, Rancurel C, Bernard T, Lombard V, Henrissat B. The Carbohydrate-Active EnZymes database (CAZy): an expert resource for glycogenomics. Nucleic Acids Res 2009; 37: D233–D238.

[70] Brulc JM, Antonopoulos DA, Berg Miller ME, Wilson MK, Yannarell AC, Dinsdale EA, et al. Gene-centric metagenomics of the fiber-adherent bovine rumen microbiome reveals forage specific glycoside hydrolases. Proc Natl Acad Sci 2009; 106: 1948–1953.

[71] Patel DD, Patel AK, Parmar NR, Shah TM, Patel JB, Pandya PR, et al. Microbial and carbohydrate active enzyme profile of buffalo rumen metagenome and their alteration in response to variation in the diet. Gene 2014; 545: 88–94.

[72] Stewart CS, Flint HJ, Bryant MP. The rumen bacteria. The rumen microbial ecosystem. 1997. Springer, pp 10–72.

[73] Vértessy BG, Tóth J. Keeping uracil out of DNA: physiological role, structure and catalytic mechanism of dUTPases. Acc Chem Res 2009; 42: 97–106.

[74] Shkoporov AN, Clooney AG, Sutton TDS, Ryan FJ, Daly KM, Nolan JA, et al. The human gut virome is highly diverse, stable, and individual specific. Cell Host Microbe 2019; 26: 527–541.

[75] Van Espen L, Bak EG, Beller L, Close L, Deboutte W, Juel HB, et al. A previously undescribed highly prevalent phage identified in a Danish enteric virome catalog. mSystems 2021; 6: e00382–21.

[76] Minot S, Sinha R, Chen J, Li H, Keilbaugh SA, Wu GD, et al. The human gut virome: inter-individual variation and dynamic response to diet. Genome Res 2011; 21: 1616–1625.

[77] Leahy SC, Kelly WJ, Altermann E, Ronimus RS, Yeoman CJ, Pacheco DM, et al. The genome sequence of the rumen methanogen *Methanobrevibacter ruminantium* reveals new possibilities for controlling ruminant methane emissions. PLoS One 2010; 5: e8926.

[78] Cottle DJ, Nolan J V, Wiedemann SG. Ruminant enteric methane mitigation: a review. Anim Prod Sci 2011; 51: 491–514.

[79] Lobo RR, Faciola AP. Ruminal phages–a review. Front Microbiol 2021; 12: 763416.

[80] Altermann E, Reilly K, Young W, Ronimus RS, Muetzel S. Tailored nanoparticles with the potential to reduce ruminant methane emissions. Front Microbiol 2022; 13: 816695.

